# ABA-dependent and ABA-independent functions of RCAR5/PYL11 in response to cold stress

**DOI:** 10.1101/723627

**Authors:** Chae Woo Lim, Sung Chul Lee

**Affiliations:** Department of Life Science, Chung-Ang University, Seoul 06974, Republic of Korea

**Keywords:** abscisic acid, cold stress, RCAR, seed germination, stomatal closure

## Abstract

*Arabidopsis thaliana* has 14 abscisic acid (ABA) receptors—PYR1/PYLs/RCARs—which have diverse and redundant functions in ABA signaling; however, the precise role of these ABA receptors remains to be elucidated. Here, we report the functional characterization of RCAR5/PYL11 in response to cold stress. Expression of *RCAR5* gene in dry seeds and leaves was ABA-dependent and ABA-independent, respectively. Under cold stress conditions, seed germination was markedly delayed in *RCAR5*-overexpressing (*Pro35S:RCAR5)* plants, but not in *Pro35S:RCAR5* in ABA-deficient (*aba1-6*) mutant background. Leaves of *Pro35S:RCAR5* plants showed enhanced stomatal closure—independent of ABA—and high expression levels of cold, dehydration, and/or ABA-responsive genes; these traits conferred enhanced freezing tolerance. Our data suggest that RCAR5 functions in response to cold stress by delaying seed germination and inducing rapid stomatal closure via ABA-dependent and ABA-independent pathways, respectively.

## Introduction

The plant hormone abscisic acid (ABA) plays a key role in growth and development, as well as in adaptive mechanisms to unfavorable conditions such as regulation of seed dormancy, germination, and stomatal opening and closure (Cutler *et al.*, 2010, Hubbard *et al.*, 2010). During seed maturation, ABA accumulates in seeds, inducing and maintaining seed dormancy and inhibiting seed germination by preventing water uptake and endosperm rupture (Müller *et al.*, 2006, Seo *et al.*, 2006). Seed germination leads to a rapid decline in ABA content and suppression of ABA signaling, suggesting that ABA is an important hormone inhibiting seed germination until favorable growth conditions prevail (Finkelstein *et al.*, 2008, Shu *et al.*, 2016). In vegetative tissue, drought stress induces ABA accumulation, leading to stomatal closure and induction of stress-related genes (Cutler *et al.*, 2010). In Arabidopsis, ABA levels increase transiently and less markedly (2-fold) in response to cold stress than in response to drought stress (approx. 20-fold) (Lång *et al.*, 1994). Several studies have shown that ABA is involved in cold stress responses; these studies have examined cold stress-induced ABA biosynthesis (Cuevas *et al.*, 2008, Lång *et al.*, 1994); absence of cold acclimation in ABA-deficient mutants (Gilmour & Thomashow, 1991, Xiong *et al.*, 2001); and ABA induction of cold-responsive genes, mainly CBF genes (Knight *et al.*, 2004, Lee & Seo, 2015). However, in comparison with the well documented role of ABA in drought stress, the function of this plant hormone in cold stress and acclimation remains controversial.

ABA is perceived by the PYRABACTIN RESISTANCE/PYRABACTIN RESISTANCE-LIKE/REGULATORY COMPONENT OF ABA RECEPTOR (PYR/PYL/RCAR; hereafter referred to as RCARs) protein family, which consists of 14 members in Arabidopsis. RCARs belong to the START-domain superfamily and are divided into three subfamily groups according to sequence homology (Ma *et al.*, 2009, Park *et al.*, 2009). In response to environmental stresses, ABA is rapidly synthesized and binds to RCARs (Cutler *et al.*, 2010). ABA-bound RCARs selectively interact with and inhibit clade A protein phosphatase 2Cs (PP2Cs), including ABA-insensitive 1 (ABI1), ABI2, hypersensitive to ABA1 (HAB1), HAB2, ABA-hypersensitive germination 1 (AHG1), AHG3/PP2CA, highly ABA-induced PP2C gene 1 (HAI1), HAI2, and HAI3 (Bhaskara *et al.*, 2012, Gonzalez-Guzman *et al.*, 2012, Ma *et al.*, 2009, Nishimura *et al.*, 2010, Park *et al.*, 2009). Among potential RCAR–PP2C interactions, 113 pairings are known to be functional (Tischer *et al.*, 2017). As a consequence of these interactions, PP2C inhibition of sucrose non-fermenting 1-related protein kinase 2s (SnRK2s) is cancelled, resulting in phosphorylation and activation of downstream components, such as transcription factors and ion channels (Furihata *et al.*, 2006, Geiger *et al.*, 2009, Lee *et al.*, 2009). In this process, bZIP transcription factor ABI5—the core ABA signaling component— is phosphorylated by ABA-activated SnRK2.2, SnRK2.3, and SnRK2.6 (Nakashima *et al.*, 2009) and subsequently regulates expression of stress adaptation genes, e.g., late embryonic and abundant genes such as *EARLY METHIONINE-LABELED 1* (*EM1*) and *EM6* (Finkelstein & Lynch, 2000).

RCARs have diverse and redundant functions in ABA and drought-stress signaling (Antoni *et al.*, 2013, Hao *et al.*, 2011, Zhao *et al.*, 2016, Zhao *et al.*, 2013). With the exception of RCAR7, RCARs induce expression of ABA-responsive genes with highly variable expression levels. Also, RCAR levels—with the exception of RCAR4—significantly suppressed by ABA deficiency (Fujii *et al.*, 2009, Ma *et al.*, 2009, Szostkiewicz *et al.*, 2010, Tischer *et al.*, 2017, Zhao *et al.*, 2013). This may be explained by differing ABA affinities and ABA dependencies of RCARs and/or variations in abundance of RCARs and their target PP2Cs (Tischer *et al.*, 2017). ABA affinities of RCARs are influenced by oligomerization states; monomeric RCARs (RCAR1, RCAR3, RCAR4, RCAR8, RCAR9, and RCAR10) have higher affinities than dimeric RCARs (RCAR11, RCAR12, RACR13, and RCAR14) (Dupeux *et al.*, 2011, Hao *et al.*, 2011). Moreover, monomeric RCARs interact with PP2Cs mainly in an ABA-independent manner, whereas dimeric RCARs bind to PP2Cs in an ABA-dependent manner (Hao *et al.*, 2011, Tischer *et al.*, 2017). In most RCARs, functional redundancy interferes with analysis of biological function in ABA signaling based on single-gene mutations (Park *et al.*, 2009). In contrast, mutation in multiple *RCARs* negatively influences ABA sensitivity and drought resistance, whereas overexpression of *RCARs* positively regulates ABA signaling and drought response (Gonzalez-Guzman *et al.*, 2012, Lim *et al.*, 2014, Santiago *et al.*, 2009, Zhao *et al.*, 2014). Loss-of-function of *RCAR3/PYL8* resulted in reduced sensitivity to ABA-induced inhibition of primary root growth (Antoni *et al.*, 2013). In combination with *RCAR2/PYL9* and *RCAR3/PYL8* functions in regulating of lateral root growth via interaction with MYB transcription factor MYB77 and MYB44 by independent manner of the core ABA signaling pathway (Xing *et al.*, 2016, Zhao *et al.*, 2014). *RCAR2/PYL9* overexpression conferred drought resistance by promoting ABA-mediated leaf senescence (Zhao *et al.*, 2016). This suggests that RCARs have distinct functions, which are associated with different tissues, developmental stages, and specific environmental conditions (Sun *et al.*, 2017). However, non-redundant functionality of RCAR in an ABA-dependent and/or ABA-independent manner remains to be elucidated. Here, we attempted to functionally characterize RCAR5 in response to cold stress.

## Results

### *RCARs* are differentially regulated in ABA-deficient seeds

Several studies based on public microarray data have shown that expression patterns of *RCARs* vary among different tissues and in response to ABA and abiotic stresses, suggesting substantial functional differences among these genes (Gonzalez-Guzman *et al.*, 2012, Kilian *et al.*, 2007, Klepikova *et al.*, 2016, Park *et al.*, 2009, Santiago *et al.*, 2009, Szostkiewicz *et al.*, 2010, Winter *et al.*, 2007). However, *RCAR* gene expression patterns in seeds remain unclear. We investigated expression levels of *RCARs* in dry seeds of *Arabidopsis thaliana* Columbia-0 (Col-0) and Landsberg *erecta* (Ler) ecotypes (Appendix Fig S1A). Transcripts of six *RCARs*—*RCAR1, RCAR2, RCAR3, RCAR5, RCAR6*, and *RCAR11*—were relatively abundant; further, *RCAR1, RCAR2, RCAR5*, and *RCAR6* were significantly downregulated after imbibition (Appendix Fig S1B and C). ABA is a key hormone involved in the induction and maintenance of seed dormancy, and its level decreases rapidly following imbibition (Kushiro *et al.*, 2004, Millar *et al.*, 2006). Hence, we postulated that transcriptional alteration of *RCAR1, RCAR2, RCAR5*, and *RCAR6* is associated with endogenous ABA level in seeds. We examined expression levels of *RCARs* in the ABA-deficient mutants *aba1-6* (Col-0 background) and *aba1-3* (Ler background). In comparison with wild-type (WT) seeds, ABA-deficient seeds showed upregulated expression of *RCAR9* and *RCAR10*, but downregulated expression of *RCAR1, RCAR2, RCAR5*, and *RCAR6* (Fig 1A). Hence, *RCAR* expression in seeds may be influenced by endogenous ABA level, and downregulation of *RCAR1, RCAR2, RCAR5*, and *RCAR6* may be associated with breaking of ABA-induced seed dormancy.

**Figure 1.**
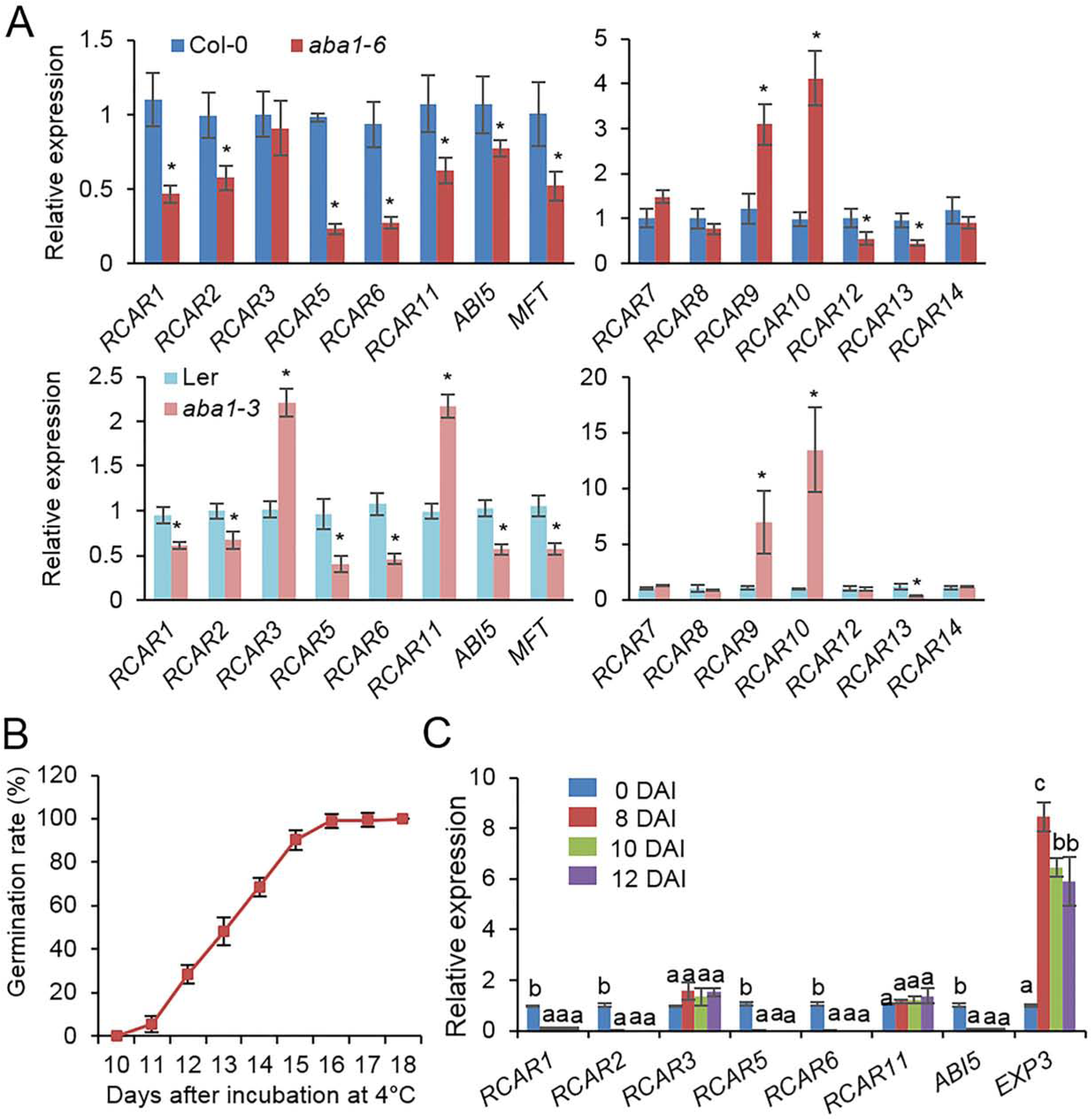
*RCAR* expression in Arabidopsis dry seeds and during seed germination under cold stress conditions. A Differential *RCAR* expression in dry seeds of ABA-deficient mutants *aba1-6* (Col-0 background; upper) and *aba1-3* (Ler background; bottom). *PP2A* was used as an internal control for normalization. The expression level of each *RCAR* in wild-type (WT) Arabidopsis seeds was set to 1.0. ABA-responsive gene *ABI5* and MFT were used as positive controls. B Germination rate of WT (Col-0) seeds under cold stress (4°C) conditions. For germination rate, seeds with radicle protrusion were counted as germinated. C *RCAR* expression during seed germination under cold stress conditions. ABA-responsive gene *ABI5* and cell wall-related gene *EXP3* were used as controls for seed germination. All data represent mean ± standard deviation (SD) of three independent experiments. Asterisks and different letters indicate significant difference between samples (Student’s *t*-test for A and ANOVA test for C, *P* < 0.05).

Dormancy is an important adaptive trait that improves plant survival by inhibiting seed germination under unfavorable conditions, including cold stress (Finkelstein *et al.*, 2008). In general, cold stratification (2–5°C for 2–4 days) causes a release of seed dormancy in Arabidopsis and promotes seed germination (Baskin & Baskin, 2004). We sowed freshly harvested seeds of Arabidopsis Col-0 on 0.5× MS agar plates, stratified the seeds at 4 ± 1°C in the dark for 2 days, and subjected the stratified seeds to cold stress by continuous incubation at 4 ± 1°C. Under cold stress conditions, germination was delayed; seeds started to germinate 10–11 days after incubation (DAI) and all seeds had germinated 16 DAI (Fig 1B). We investigated the functional role of *RCARs* in sensitivity of imbibed seeds to cold stress. First, we examined the expression patterns of *RCARs* during seed germination under cold stress conditions (Fig 1C). Genes involved in ABA response and GA-mediated cell wall modification are differentially expressed during seed germination (Liu *et al.*, 2016). We amplified the ABA-responsive gene *ABI5* and the cell wall-related gene *EXPANSIN3* (*EXP3*) as a control; at 8 DAI, transcripts of *ABI5* and *EXP3* were significantly downregulated and upregulated, respectively. Consistent with data obtained using *Arabidopsis thaliana* Col-0 and Ler ecotypes (Appendix Fig S1B and C), expression levels of *RCAR1, RCAR2, RCAR5*, and *RCAR6* decreased significantly during seed germination; however, we determined no significant change in expression levels of *RCAR3* and *RCAR11* (Fig 1C).

### RCAR5 plays a key role in delay of seed germination under cold stress conditions

Several studies have shown that ABA signaling is related to the cold stress response (Ding *et al.*, 2015, Knight *et al.*, 2004, Kreps *et al.*, 2002, Lee *et al.*, 2010). To analyze the functional role of *RCARs* in seed germination under cold stress conditions, we generated Arabidopsis transgenic plants overexpressing each *RCAR* under the control of the 35S promoter. In comparison with WT plants, the 14 transgenic lines showed high expression levels of each *RCAR* gene (>30-fold to 300-fold) (Appendix Fig S1D) and also high sensitivity to ABA during seed germination and seedling growth (Fig 2A). To examine the mechanism by which enhanced ABA sensitivity influences seed germination or seedling growth of transgenic plants under cold stress conditions, freshly harvested seeds of each line were sown on 0.5× MS agar plates and stratified at 4 ± 1°C in the dark for 2 days. Under normal growth conditions (24°C), germination rates did not differ significantly between transgenic plants and WT plants (Fig 2A and Appendix Fig S1E–G). However, under cold stress conditions, the germination rates of transgenic lines *Pro35S:RCAR2 and Pro35S:RCAR5* at 14 DAI were significantly lower (56.0% and 25.9%, respectively) than those of WT plants (64.4%) (Fig 2B). At 21 DAI, the number of etiolated seedlings was significantly lower in *Pro35S:RCAR5* lines than in WT and *Pro35S:RCAR2* lines but did not differ significantly between WT and *Pro35S:RCAR2* lines. We further observed phenotypic differences in seedling growth at 8°C; at this temperature, germination of WT and transgenic lines was enhanced; however, the germination rate was significantly lower in *Pro35S:RCAR5* lines than in WT and the remaining 13 transgenic lines (Fig 2A). To determine whether there was a loss of seed viability in the *Pro35S:RCAR5* line, seeds of *Pro35S:RCAR5* (#1) and *Pro35S:RCAR5* (*#2*) were incubated at 4 ± 1°C for 18 days and maintained at 24°C. After 4 days, all seeds germinated and developed into seedlings (Fig 2C).

**Figure 2.**
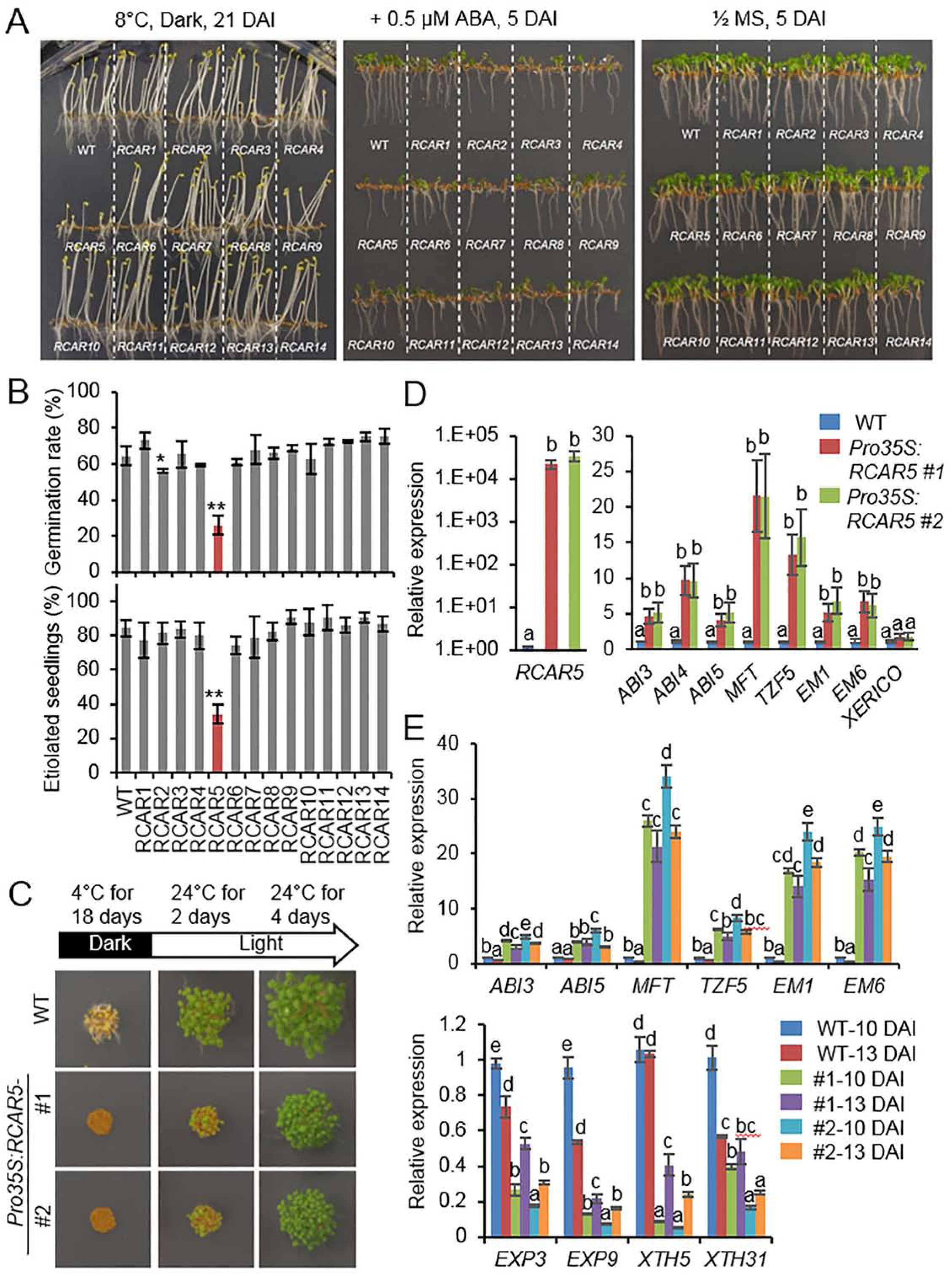
Delayed germination of *Pro35S:RCAR5* seeds under cold stress conditions. A Seedling development of *Pro35S:RCAR* transgenic lines and WT plants exposed to cold stress and ABA. Seeds of *Pro35S:RCAR* transgenic lines were germinated on 0.5× MS medium supplemented with 0 μM or 0.5 μM ABA and vertically grown at 8°C in the dark (left) and at 24°C in the light (middle and right). Representative images were taken at the indicated time points. DAI, days after incubation. B Germination and seedling growth of *Pro35S:RCAR* mutants under cold stress conditions. For measuring germination rates and seedling growth of each plant line, the numbers of germinated seeds (with an emerged radicle) and etiolated seedlings were counted at 13 DAI and 21 DAI, respectively. C Recovery of delayed germination of *Pro35S:RCAR5* seeds under normal growth conditions. After incubation for 18 days at 4°C in the dark, *Pro35S:RCAR5* seeds were transferred to 24°C with light. Representative images were taken at the indicated time points. D Expression patterns of ABA-responsive genes in dry seeds of *Pro35S:RCAR* mutants. E Expression patterns of ABA-responsive genes and cell-wall related genes in seeds of *Pro35S:RCAR* mutants and WT plants during germination at 4°C in the dark. All data represent mean ± SD of three independent experiments. Asterisks and different letters indicate significant difference compared with WT (Student’s *t*-test for B and ANOVA test for D and E; \**P* < 0.05; \*\**P* < 0.01).

To examine delayed germination of *Pro35S:RCAR5* seeds at the molecular level, we performed quantitative reverse transcription-polymerase chain reaction (qRT-PCR) analysis. We examined expression levels of ABA-responsive genes in dry seeds of *Pro35S:RCAR5* and WT lines (Fig 2D). The ABA-responsive genes *ABA INSENSITIVE 3* (ABI3), *ABI4, ABI5, MOTHER OF FT AND TFL1* (*MFT*), *TANDEM CCCH ZINC FINGER PROTEIN 5* (*TZF5*), *EARLY METHIONINE-LABELED 1* (*EM1*), and *EM6* were more strongly expressed in *Pro35S:RCAR5* seeds than in WT seeds. However, we determined no significant difference in expression level of the *XERICO* gene—which promotes ABA accumulation in seeds (Zentella *et al.*, 2007)—between *Pro35S:RCAR5* and WT seeds. During germination at 4°C, expression levels of ABA-responsive genes were significantly higher in *Pro35S:RCAR5* seeds than in WT seeds; however, expression of cell wall-related genes—*EXPANSIN3* (*EXP3*), *EXP9, XYLOGLUCAN ENDOTRANSGLUCOSYLASE/HYDROLASE* (*XTH5*), and *XTH31*—was downregulated in *Pro35S:RCAR5* seeds (Fig 2E). Our data suggest that *RCAR5* functions in ABA-mediated inhibition of seed germination under cold stress conditions.

### Cold stress-induced germination delay of *Pro35S:RCAR5* seeds occurs via an ABA-dependent pathway

*Pro35S:RCAR5* seeds showed high expression levels of ABA-responsive genes; hence, we postulated that *RCAR5* overexpression delays seed germination via an ABA-dependent pathway. WT and *Pro35S:RCAR5* seeds were placed on 0.5× MS medium supplemented with norflurazon (NF)—an ABA biosynthesis inhibitor—and germinated at 4°C in the dark. Supplementation with NF increased the germination rate in a dose-dependent manner; notably, at 14 DAI, the germination rate of *Pro35S:RCAR5* seeds was 2-6 fold higher in the presence than in the absence of NF (Fig 3A and B). This enhanced germination was observed at all the investigated time points. In contrast, at 16 DAI, we observed no influence of NF on germination of WT seeds. In contrast to NF, ABA strongly inhibited seed germination and caused a 4-day delay in germination of WT seeds (Fig 3C). At 18 DAI, the germination rates of WT and *Pro35S:RCAR5* seeds on 0.5× MS medium supplemented with 0.75 μM ABA were 77.1% and <3%, respectively. As predicted, NF was antagonistic to ABA-mediated inhibition of seed germination. In the presence of NF, the germination rate of WT seeds at 14 DAI was 48.2–64.5% (vs. 5.6% in the absence of NF); moreover, in the presence of ABA, NF enhanced germination of *Pro35S:RCAR5* seeds by up to 26%. To verify germination delay of *Pro35S:RCAR5* seeds using genetic analysis, we overexpressed *RCAR5* in the ABA-deficient mutant *aba1-6* (*Pro35S:RCAR5/aba1-6*; Appendix Fig S2A) and analyzed the relationship between RCAR5-mediated ABA signaling and delayed seed germination under cold stress conditions. At 4°C, the germination rate of *aba1-6* seeds was 1.3–1.9-fold higher than that of WT seeds; moreover, germinated *aba1-6* seedlings had longer radicles than WT seedlings (Fig 3D and E). *RCAR5* overexpression did not alter the germination rate of *aba1-6* mutants at 4°C. However, similar to *Pro35S:RCAR5* seedlings, *Pro35S:RCAR5/aba1-6* seedlings were more sensitive to ABA—as measured by lower seed germination rates and shorter radicles—than were *aba1-6* seedlings (Appendix Fig S2B–E). Owing to this ABA hypersensitivity, application of exogenous ABA strongly inhibited germination of *Pro35S:RCAR5/aba1-6* seeds under cold stress conditions (Fig 3F). Our data suggest that RCAR5 contributes to cold stress-induced delay of seed germination via an ABA-dependent pathway.

**Figure 3.**
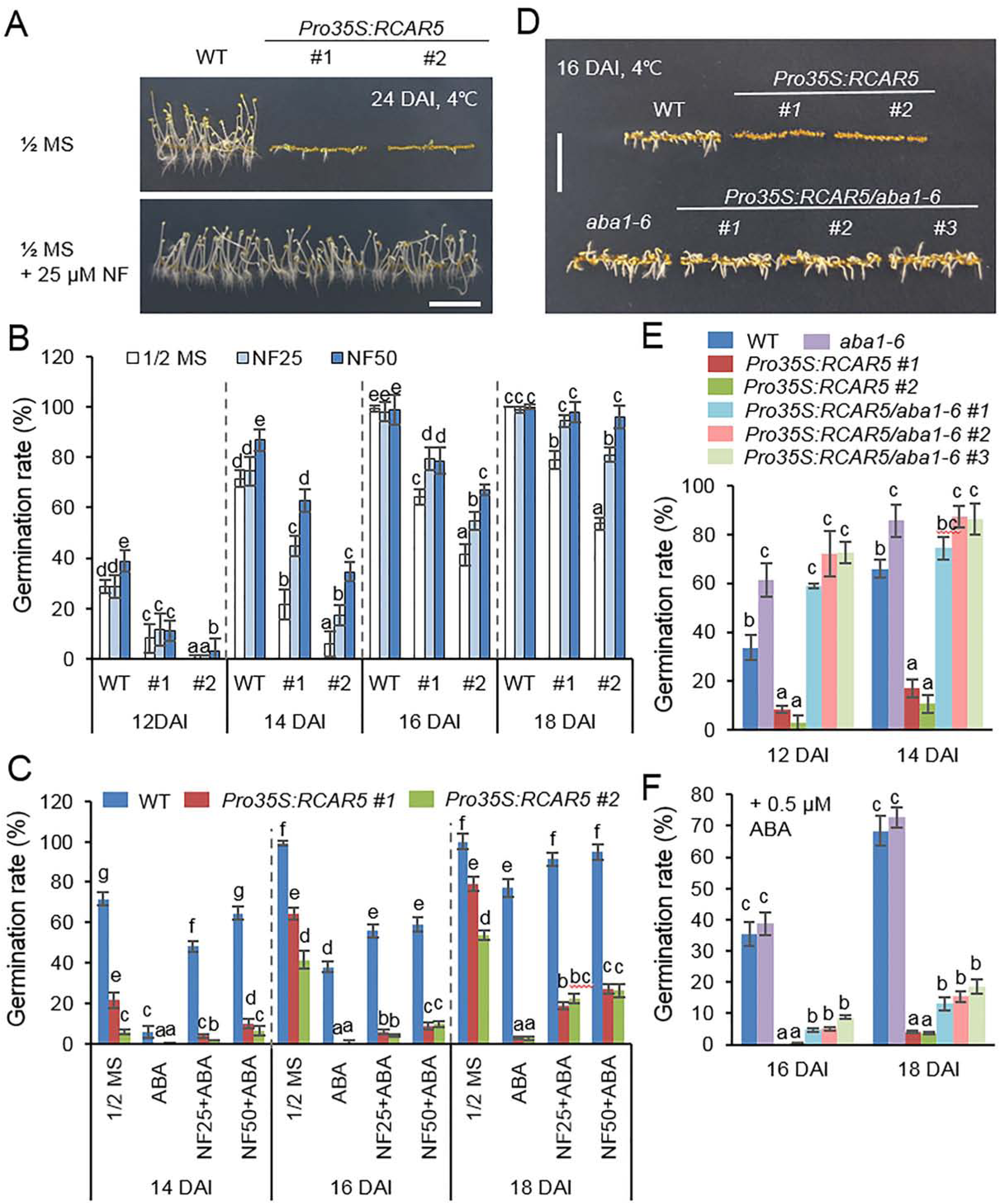
Cold-induced germination delay of *Pro35S:RCAR5* seeds in an ABA-dependent manner. A-C Effect of norflurazon on seed germination of WT and *Pro35S:RCAR5* lines. Seeds of *Pro35S:RCAR5* and WT were germinated on 0.5× MS medium supplemented with norflurazon (25 μM or 50 μM) and/or ABA (0.75 μM) and vertically grown at 4°C in the dark. After 24 days, representative images were taken (A). For measuring germination rates of each plant line, the numbers of seeds with emerged radicles were counted at the indicated time points (B and C). D-F Germination and seedling growth of *Pro35S:RCAR/aba1-6* mutants under cold stress conditions. Seeds of each transgenic line were plated on 0.5× MS medium supplemented with 0 μM or 0.75 μM ABA and germinated in the dark at 4°C. Representative images were taken at 16 DAI (D) and germination rate was measured at the indicated time points (E and F). All data represent mean ± SD of three independent experiments. Different letters indicate significant difference compared with WT (ANOVA; *P* < 0.05). Scale bar = 1 cm.

### HAB1, but not ABI5, is involved in in delayed germination of *Pro35S:RCAR5* seeds under cold stress conditions

To examine the mechanism whereby germination of *Pro35S:RCAR5* seeds is delayed under cold stress conditions, we genetically analyzed the downstream components of RCAR5 in ABA signaling. Yeast two-hybrid and bimolecular fluorescence complementation assay (BiFC) analyses revealed the interaction of RCAR5 with HAB1 and PP2CA (Appendix Fig S3A and B); RCAR5–HAB1 interaction occurred in the nucleus and cytoplasm, whereas RCAR5– PP2CA interaction occurred only in the nucleus. The expression levels of *HAB1* and *PP2CA* in seeds decreased after stratification or imbibition during germination (Appendix Fig S4A and B) and increased in cold stress-treated seeding (Appendix Fig S4C-D). Loss-of-function mutants of *HAB1* and *PP2CA* showed ABA hypersensitivity during germination and seedling growth (Appendix Fig S3C and D) (Lim *et al.*, 2014, Saez *et al.*, 2004a). Consistently, under cold stress conditions, seed germination and seedling growth were more strongly inhibited in *hab1* and *pp2ca* mutants than in WT plants; however, at 16 DAI, the germination rates of *hab1* and *pp2ca* mutants were similar to those of WT plants (Appendix Fig S3C and D). We previously reported that HAB1 acts downstream of RCAR5 in ABA signaling (Lim & Lee, 2015); hence, we examined the functional relationship between *HAB1* and *RCAR5* during seed germination at 4°C. In contrast to *hab1* seeds, *Pro35S:HAB1* seeds germinated more rapidly than WT seeds (Fig 4A and B). At 13 DAI, the germination rate of *Pro35S:HAB1/Pro35S:RCAR5* seeds did not differ significantly from that of *Pro35S:RCAR5* seeds; thereafter, the germination rate of *Pro35S:HAB1/Pro35S:RCAR5* seeds was significantly higher than that of *Pro35S:RCAR5* seeds. Our results indicate that *HAB1* overexpression compromises RCAR5-induced ABA hypersensitivity in *Pro35S:HAB1/Pro35S:RCAR5* seeds; these findings are in agreement with our ABA hyposensitivity data (Appendix Fig S4E-I).

**Figure 4.**
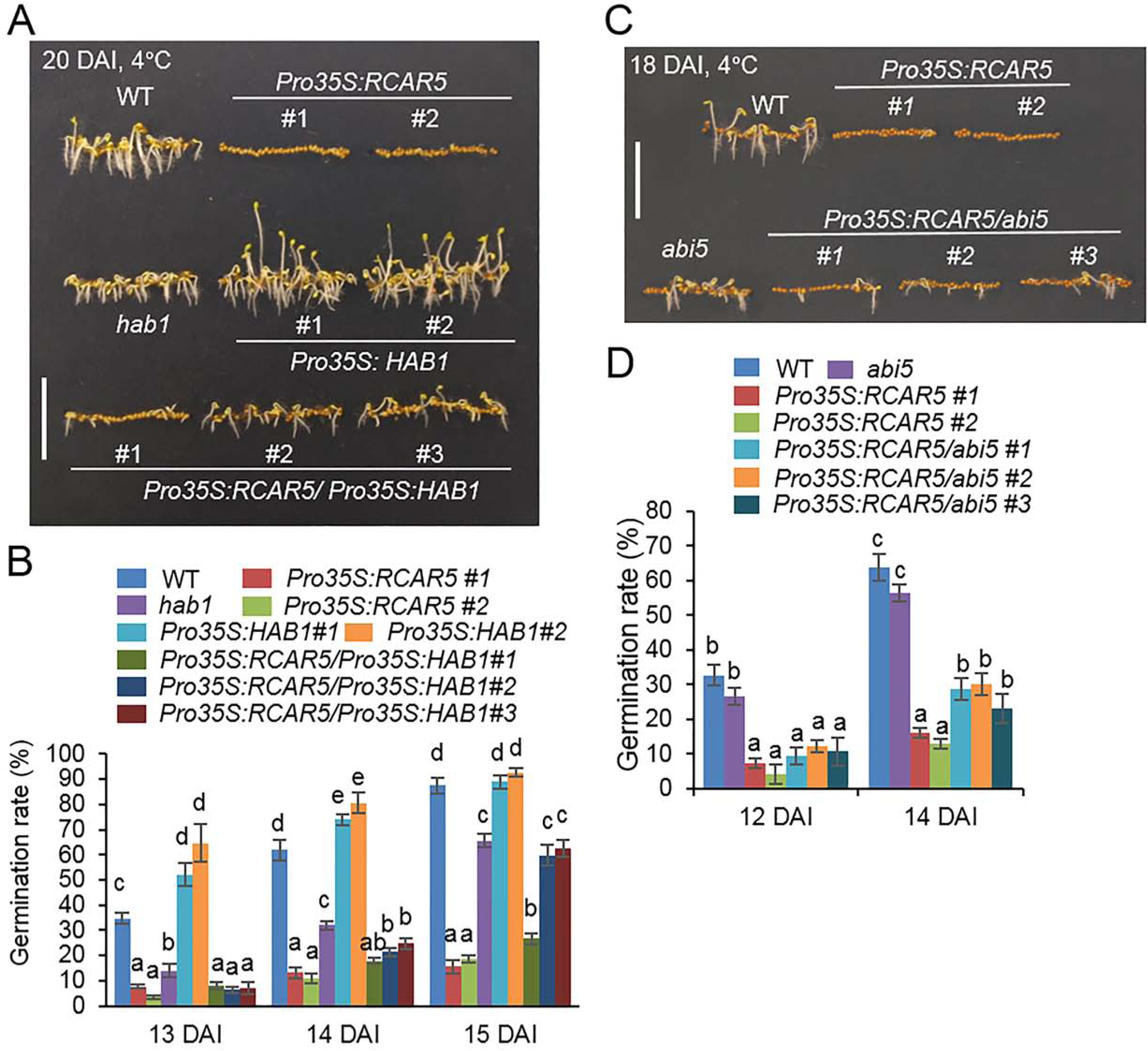
Delayed germination of *Pro35S:RCAR5* seeds under cold stress conditions is regulated by downstream component HAB1, but not by ABI5. A, B Germination rates of *Pro35S:RCAR5, hab1, Pro35S:HAB1, Pro35S:RCAR5/Pro35S:HAB1*, and WT plants under cold stress conditions. Seeds were plated on 0.5× MS medium and germinated in the dark at 4°C. For measuring germination rates, the numbers of seeds with emerged radicles were counted at the indicated time points (B). Representative images were taken at 20 DAI (A). C, D Germination rates of *Pro35S:RCAR5, abi5, Pro35S:RCAR5/abi5*, and WT plants under cold stress conditions. Representative images were taken at 18 DAI (C) and germination rates were determined at 12 DAI and 14 DAI (D). Data represent mean ± SD of three independent experiments, each evaluating 100 seeds. Different letters indicate significant difference compared with WT (ANOVA; *P* < 0.05). Scale bar = 1 cm.

Next, we used the ABA-insensitive—*abi5* [designated as *abi5-8* by Zheng *et al.* (2012b)]—mutant. As a bZIP transcription factor, ABI5 is mainly expressed in dry seeds and plays a positive role in ABA signaling during seed germination and early seedling growth (Finkelstein & Lynch, 2000, Lopez-Molina *et al.*, 2001). *abi5* mutants showed ABA hyposensitivity (Zheng *et al.*, 2012b), whereas *ABI5*-overexpressing plants were hypersensitive to ABA (Lopez-Molina *et al.*, 2001). We overexpressed *RCAR5* in *abi5* (*Pro35S:RCAR5/abi5*) mutants; ABA sensitivity—as measured by germination rate and seedling establishment—of these mutants was similar to that of *abi5* mutants (Appendix Fig S2F-J). Based on the observed ABA hyposensitivity, we predicted increased germination of *abi5* and *Pro35S:RCAR5/abi5* seeds under cold stress conditions. However, the germination rate of *abi5* seeds was lower than that of WT seeds, although the difference was not significant; moreover, at 18 DAI, *abi5* seedlings were smaller than WT seedlings (Fig 4C and D). *Pro35S:RCAR5/abi5* seed germination at 4°C was markedly delayed; however, germination rates were significantly higher than those of *Pro35S:RCAR5* lines and significantly lower than those of WT seeds (Fig 4D). Our data suggest that cold stress-induced delay in the germination of the *Pro35S:RCAR5* lines is regulated by HAB1, but not by ABI5.

### *RCAR5* is induced in leaves in response to cold stress in an ABA-independent manner

To investigate the functional role of RCARs in leaves in response to cold stress, we analyzed expression patterns of *RCARs* in Arabidopsis leaves in response to cold stress (4°C). In the eFP browser (Winter *et al.*, 2007), several *RCARs* were downregulated by cold stress; however, no data on *RCAR4, RCAR5, RCAR6*, or *RCAR7* that have much lower expression levels in the leaves relative to the other *RCARs* were available (Appendix Fig S5A-C). qRT-PCR analysis of 14 *RCARs* revealed cold stress-induced alteration of *RCAR* gene expression. Consistent with our in-silico data, expression levels of *RCAR7, RCAR8, RCAR10, RCAR11, RCAR12*, and *RCAR13* gradually decreased in response to cold stress, whereas the expression levels of *RCAR1, RCAR2, RCAR5*, and *RCAR6* were strongly induced (>2-fold) relative to the control (Fig 5A). *RCAR5* showed the highest induction level (up to 4-fold). ABA deficiency did not alter the expression level of *RCAR5* in WT leaves (Appendix Fig S5D). To examine whether cold stress-induced *RCAR5* expression occurs in an ABA-independent manner, we analyzed the expression pattern of *RCAR5* in the ABA-deficient mutants *aba1-6* and *aba1-3* (Ler background) and the ABA-insensitive mutant *abi5* under cold stress conditions. Similar to data obtained using the Col-0 ecotype (Fig 5A), the investigated mutants showed gradual expression of *RCAR5* in rosette leaves in response to cold stress (Fig 5B), indicating that *RCAR5* is induced by cold stress via an ABA-independent pathway.

**Figure 5.**
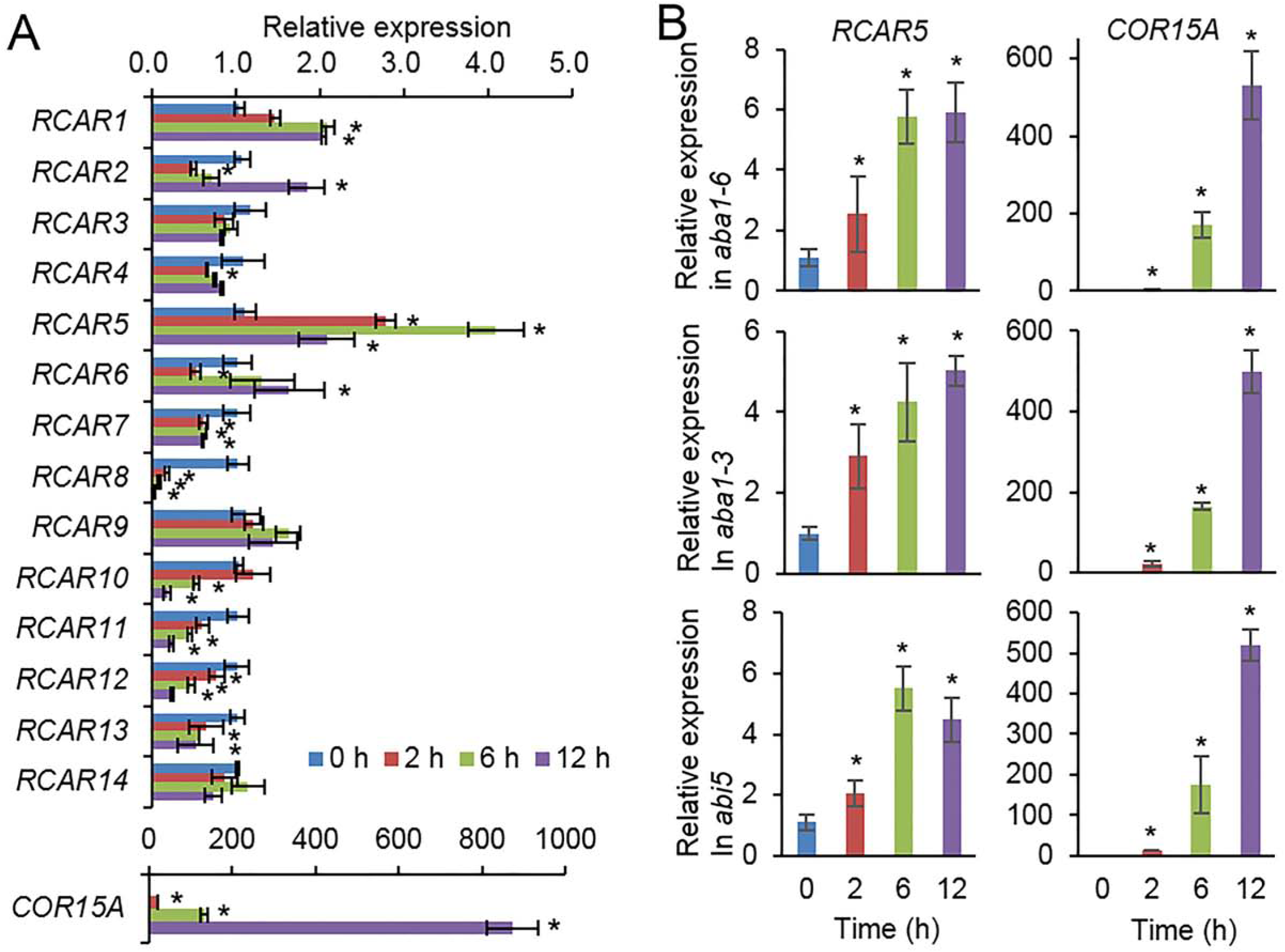
ABA-independent *RCAR5* expression in Arabidopsis leaves in response to LT. A *RCAR* expression in response to cold stress. Three-week-old Arabidopsis Col-0 plants were exposed to 4°C and rosette leaves were harvested at the indicated time points. *Actin8* was used as an internal control for normalization. *COR15A* was used as a positive control for cold stress treatment. The expression level of each gene at 0 h was set to 1.0. B *RCAR5* expression pattern in ABA-deficient mutants *aba1-6* and *aba1-3* and ABA-insensitive mutant *abi5* in response to cold stress. Data represent mean ± SD of three independent experiments. Asterisks indicate significant difference compared with non-treated control (Student’s *t*-test; *P* < 0.05).

### *Pro35S:RCAR5* transgenic plants exhibit enhanced stomatal closure in response to cold stress

Cold stress triggers leaf wilting and stomatal closure (Ruelland & Zachowski, 2010, Wilkinson *et al.*, 2001). Consistent with our qRT-PCR data (Fig 5B), *RCAR5* expression was not induced in leaves treated with ABA, but was strongly induced by dehydration (Appendix Fig S5E). Hence, *RCAR5* expression may be triggered by cold stress-induced leaf dehydration in an ABA-independent manner. In comparison with WT plants, *Pro35S:RCAR5* plants showed enhanced tolerance to dehydration stress via decreased transpirational water loss from the leaves (Appendix Fig S5F and G) and increased ABA-induced stomatal closure (Lim & Lee, 2015). We conducted phenotypic analysis of *Pro35S:RCAR5* plants in response to cold stress. After 24 h incubation at 4°C, WT plants were wilted, but there were no phenotypic differences between *Pro35S:RCAR5* plants and non-treated plants (Fig 6A). Interestingly, this phenomenon was observed in *Pro35S:RCAR5* plants, but not in *Pro35S:RCAR2* or *Pro35S:RCAR3* plants (Appendix Fig S6A-C). To monitor this cold stress-induced leaf dehydration in a quantitative manner, we measured the fresh weights of aerial parts of WT and *Pro35S:RCAR5* plants. In comparison with WT plants, *Pro35S:RCAR5* plants showed significantly lower water loss; after 24 h incubation at 4°C, the fresh weights of WT and *Pro35S:RCAR5* plants were decreased by approximately 44% and 29%, respectively (Fig 6B). To examine whether the decreased transpirational water loss exhibited by *Pro35S:RCAR5* plants is derived from rapid stomatal closure in response to cold stress, we measured leaf surface temperature and stomatal apertures. Under normal growth conditions, we observed no phenotypic differences between WT and *Pro35S:RCAR5* plants (Fig 6C-F). However, after incubation at 4°C, the leaf temperatures of *Pro35S:RCAR5* plants were higher than those of WT plants; after 4 h incubation, the difference in leaf temperature was statistically significant (Fig 6C and D, and Appendix Fig S6D and E). Consistently, the stomatal apertures of *Pro35S:RCAR5* plants were significantly smaller than those of WT plants (Fig 6E and F); in comparison with non-treated plants, the average stomatal apertures of WT and *Pro35S:RCAR5* plants were reduced by 31% and 40%, respectively. Our data suggest that *RCAR5* is involved in cold stress-induced stomatal closure.

**Figure 6.**
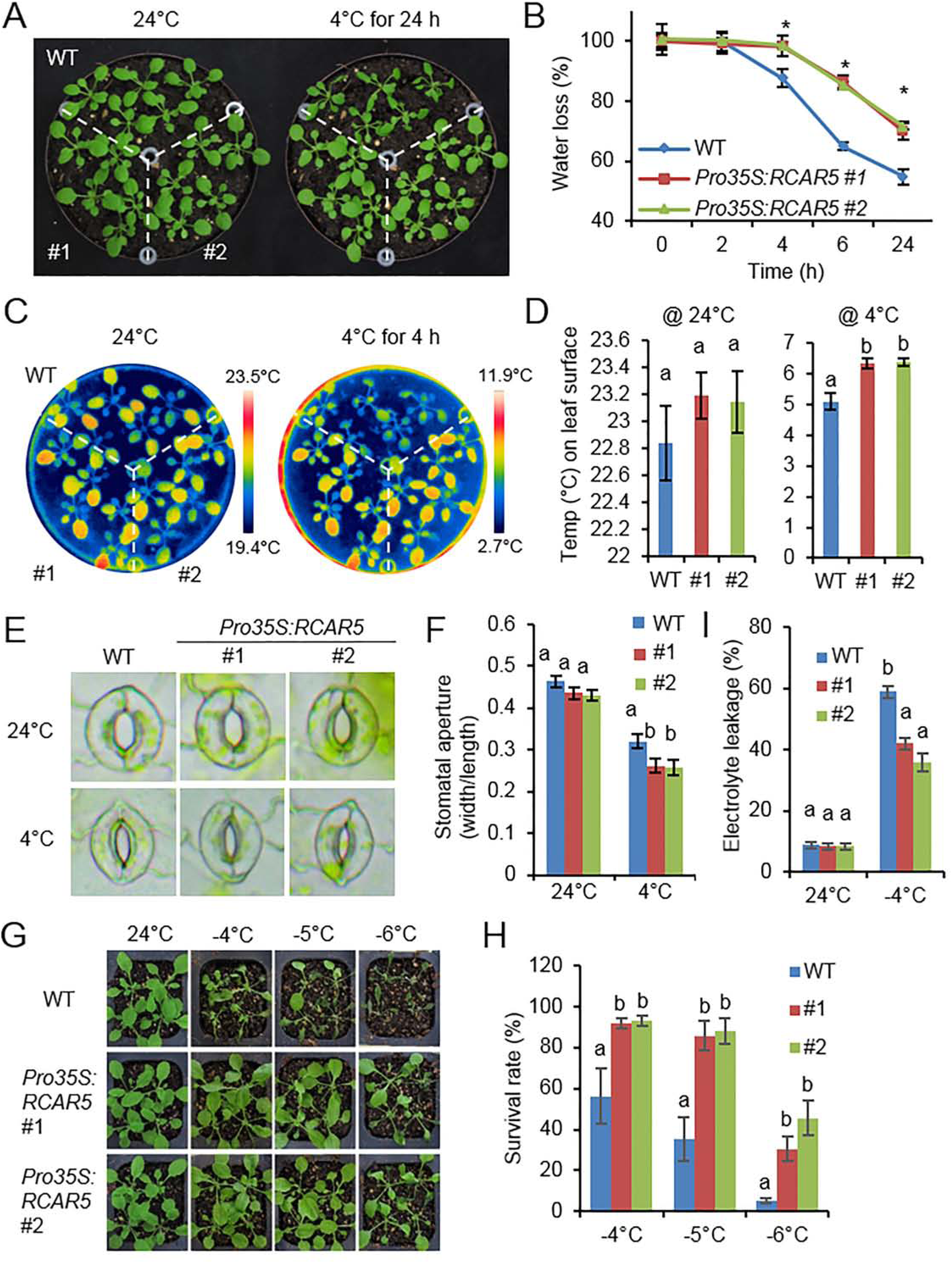
Enhanced tolerance of *Pro35S:RCAR5* plants to cold stress. A, B Cold stress-induced dehydration of *Pro35S:RCAR5* transgenic lines #1 and #2 and WT plants. Three-week-old Arabidopsis plants were treated with cold stress (4°C) for 24 h and representative images were taken (A). The fresh weights of each plant line (n = 30) were measured at the indicated time points after cold stress (B). C, D Leaf temperatures of *Pro35S:RCAR5* plants in response to cold stress. Representative thermographic images of *Pro35S:RCAR5* and WT plants 4 h after cold stress treatment (C); the mean leaf temperatures of the two largest leaves were measured using 20 plants of each line (D). E, F Stomatal apertures in *Pro35S:RCAR5* transgenic lines and WT plants treated with cold stress. Leaf peels harvested from 3-week-old plants of each line were incubated in chilled stomatal opening solution (SOS) for 2 h at 4°C. Representative images were taken (E) and the stomatal apertures were measured under the microscope (F). G, H Freezing tolerance of *Pro35S:RCAR5* and WT plants. Three-week-old seedlings of each plant line were exposed to freezing temperatures as indicated. After recovery at 24°C for 2 days, representative images were taken (G) and the survival rate of each line was counted (H). I Electrolyte leakage of *Pro35S:RCAR5* and WT plants. For freezing treatment, 3-week-old seedlings were exposed to −4°C for 1 h. All data represent mean ± SD of three independent experiments. Different letters indicate significant difference between WT and transgenic plants (ANOVA; *P* < 0.05).

### Enhanced tolerance of *Pro35S:RCAR5* transgenic plants to cold stress is accompanied by high expression levels of cold-responsive genes

*RCAR5* overexpression conferred enhanced stomatal closure in response to cold stress and ABA; hence, we postulated that *Pro35S:RCAR5* plants show enhanced tolerance to freezing stress. We exposed 3-week-old WT and *Pro35S:RCAR5* seedlings to freezing temperature (approx. −4 to −6°C) for 1 h. After transfer to 24°C for 2 days, the survival rate of *Pro35S:RCAR5* seedlings was higher than that of WT; in comparison with *Pro35S:RCAR5* seedlings, WT seedlings were severely wilted and did not survive (Fig 6G and H). Under cold stress conditions, the cellular membrane is frequently damaged because of cold stress-induced dehydration (Steponkus, 1984, Uemura *et al.*, 1995). The degree of membrane injury in plants can be evaluated by relative electrolyte leakage (Steponkus, 1984). Consistently, our *Pro35S:RCAR5* seedlings showed significantly lower electrolyte leakage than WT plants (Fig 6I).

Next, we selected 12 representative genes—*CBF1*–*3* (AT4G25490, AT4G25470, AT4G25480), *RD29A* (AT5G52310), *RD29B* (AT5G52300), *RAB18* (AT5G66400), *DREB2A* (AT5G05410), *COR15A* (AT2G42540), *COR47* (AT1G20440), *RD26* (AT4G27410), *KIN1*–*2* (AT3G21960, AT5G15970)—associated with cold stress, dehydration, and ABA signaling. We analyzed the expression patterns of these genes in response to cold stress (4°C). With the exception of *DREB2A* and *COR47*, the expression levels of the investigated genes were higher expressed in *Pro35S:RCAR5* transgenic lines than in WT plants under normal growth conditions. Moreover, the different expression levels between the two plant lines were still observed after cold stress treatment (Appendix Fig S7A). Our results suggest that high levels of CBF genes during the early stages of cold treatment may help trigger transcription of target genes in *Pro35S:RCAR5* transgenic lines. The enhanced gene expression observed in *Pro35S:RCAR5* transgenic lines may be derived from ABA hypersensitivity. *RCAR5* overexpression in ABA-insensitive *abi5* and *Pro35S:HAB1* mutant plants did not lead to a similar expression pattern in response to cold stress (Appendix Fig S7B and C). Our data indicate that upregulation of cold, dehydration, and/or ABA-induced genes may contribute to enhanced tolerance of *Pro35S:RCAR5* transgenic lines to cold stress.

### *RCAR5* overexpression in *aba1-6* mutant seedlings leads to enhanced stomatal closure in response to cold stress

ABA is a well-known phytohormone that triggers stomatal closure under abiotic stress conditions such as dehydration (Cutler *et al.*, 2010). To examine whether cold stress-induced enhanced stomatal closure in *Pro35S:RCAR5* plants is associated with ABA, we subjected 3-week-old WT, *Pro35S:RCAR5, aba1-6*, and *Pro35S:RCAR5/aba1-6* seedlings to cold stress. In comparison with WT plants, *aba1-6* and *Pro35S:RCAR5/aba1-6* mutant plants wilted rapidly. Thus, it was difficult to distinguish phenotypic differences between *Pro35S:RCAR5/aba1-6* and *aba1-6* mutant plants in response to cold stress conditions; hence, in further studies, we used only *Pro35S:RCAR5/aba1-6* and *aba1-6* plant lines. Under normal growth conditions, leaf temperatures were slightly higher in *Pro35S:RCAR5/aba1-6* than in *aba1-6* plants, but the difference was not significant; however, after cold stress treatment (4°C for 3 h), leaf temperatures were significantly higher in *Pro35S:RCAR5/aba1-6* plants than in *aba1-6* plants (Fig 7A and B). After a further 3 h of incubation, *Pro35S:RCAR5/aba1-6* plants were less wilted than *aba1-6* plants (Fig 7C). Similar to *Pro35S:RCAR5* plants, this physiological characteristic of *Pro35S:RCAR5/aba1-6* plants contributed to enhanced survival rates after freezing treatment (−4°C for 1 h) (Fig 7D). However, qRT-PCR analysis revealed that expression levels of the ABA-, dehydration-, and cold-responsive genes *RD29B, RAB18, KIN1, COR47*, and *COR15A* did not differ significantly between *Pro35S:RCAR5/aba1-6* and *aba1-6* plants in response to cold treatment (4°C for 6 h) (Fig 7E), and this may be derived from ABA deficiency. Our data suggest that cold stress induces stomatal closure via an ABA-independent pathway involving *RCAR5*.

**Figure 7.**
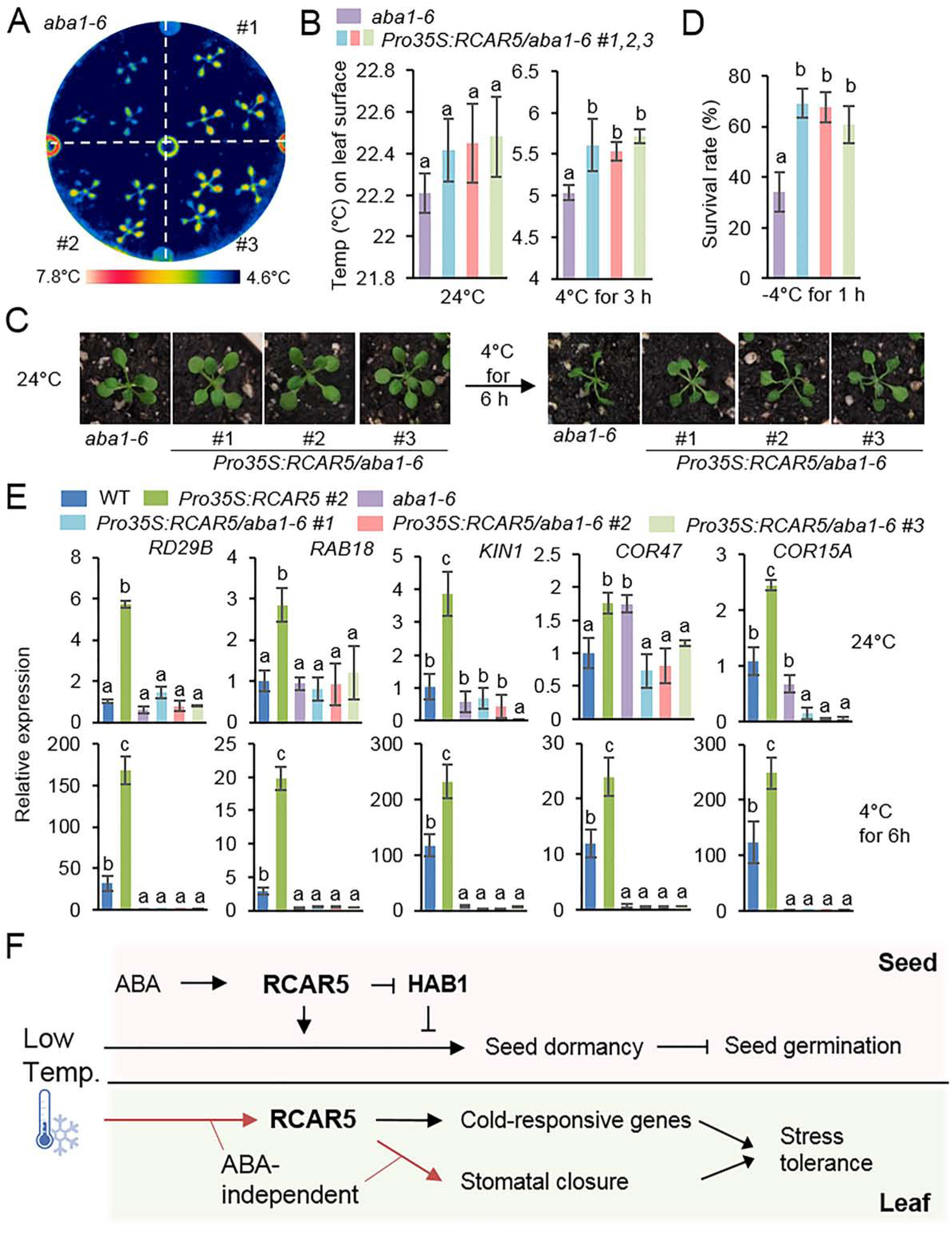
Stomatal closure and cold- and ABA-responsive gene expression of *Pro35S:RCAR5/aba1-6* plants under cold stress conditions. A, B Leaf temperatures of *Pro35S:RCAR5/aba1-6* plants in response to cold stress. Representative thermographic images of *Pro35S:RCAR5/aba1-6* and *aba1-6* plants at 3 h after cold stress treatment (A); the mean leaf temperatures of the two largest leaves were measured using 24 plants of each line (B). C Cold stress-induced dehydration of *Pro35S:RCAR5/aba1-6* and *aba1-6* plants. Three-week-old Arabidopsis plants were subjected to cold stress (4°C) for 6 h and representative images were taken. D Freezing tolerance of *Pro35S:RCAR5/aba1-6* and *aba1-6* plants. Three-week-old seedlings of each plant line were exposed to freezing temperatures (−4°C) for 1 h and the survival rate of each line was counted. E Expression patterns of cold- and ABA-responsive genes in *Pro35S:RCAR5/aba1-6* and *aba1-6* plants under cold stress conditions. F Schematic representation of the functional role of RCAR5 in ABA-mediated seed dormancy and ABA-independent stomatal closure under cold stress conditions. Arrows indicate promotion actions; lines with end bar indicate inhibitory actions.

## Discussion

As ABA receptors, Arabidopsis *RCARs* function redundantly in regulation of seed germination, stomatal aperture, and transcriptional activation in response to ABA (Gonzalez-Guzman *et al.*, 2012, Park *et al.*, 2009). Nevertheless, many studies have suggested a distinct function of RCARs in ABA and abiotic stress signaling, based on their different expression patterns, biochemical properties, and genetic analyses data (Antoni *et al.*, 2013, Dupeux *et al.*, 2011, Gonzalez-Guzman *et al.*, 2012, Hao *et al.*, 2011, Zhao *et al.*, 2016, Zhao *et al.*, 2013, Zhao *et al.*, 2014). Our data provide a new insight into the involvement of RCAR5 in cold stress responses, including delay of seed germination and rapid stomatal closure via ABA-dependent and ABA-independent pathways, respectively (Fig 7F).

Owing to functional redundancy of the *RCAR* gene family, we generated transgenic plants overexpressing each of the 14 *RCARs* that mediate ABA signaling. Overexpression of these *RCARs* conferred ABA hypersensitivity in terms of seed germination and seedling growth; however, the degree of sensitivity varied according to each *Pro35S:RCAR* mutant (Appendix Fig S1D–G). Seed dormancy and germination are influenced by ABA content and ABA sensitivity. Several studies have shown that mutations in ABA biosynthesis and signaling components influence the level of seed dormancy. Seed germination occurs more rapidly in ABA-deficient mutants—such as *aba1* and *aba2*—than in WT plants (Gonzalez-Guzman *et al.*, 2002, Xiong *et al.*, 2001). Dominant-negative mutants of *ABI1* (*abi1-1*) and *ABI2* (*abi2*) and a mutant defective in three *SnRKs* (*snrk2.2, snrk2.3*, and *snrk2.6*) showed reduced seed dormancy, consistent with their negative and positive roles in ABA signaling, respectively (Fujita *et al.*, 2009, Ma *et al.*, 2009, Nakashima *et al.*, 2009, Park *et al.*, 2009). Hence, we predicted delayed seed germination of *Pro35S:RCAR* mutants under cold stress conditions. Consistently, germination of *Pro35S:RCAR5* seeds was markedly delayed under cold stress conditions (Fig 2A and B). Germination was also delayed in *Pro35S:RCAR2* seeds, but less markedly than in *Pro35S:RCAR5* seeds. Our data suggest that the depth of seed dormancy under cold stress conditions may be only partly derived from ABA sensitivity. This discrepancy between ABA sensitivity and seed dormancy was also observed in the loss-of-function mutant of *ABI5* (*abi5*; Fig 4C and D, and Appendix Fig S2F-J). ABI5 does not control seed dormancy, but does regulate seed germination and seedling growth in response to ABA (Finkelstein & Lynch, 2000, Lopez-Molina *et al.*, 2001). Consistently, the germination rate of *abi5* seeds at 4°C did not differ significantly from that of WT seeds; moreover, similar to *Pro35S:RCAR5* seeds, germination of *Pro35S:RCAR5/abi5* seeds at 4°C was markedly delayed by *RCAR5* overexpression. However, these data do not imply that *Pro35S:RCAR5* seeds can be slowly germinated through an ABA-independent pathway. Germination of *Pro35S:RCAR5* seeds under cold stress conditions was dependent on endogenous ABA. *RCAR5*-mediated seed dormancy was blocked by ABA deficiency derived from NF treatment and loss-of-function of *ABA1* (Fig 3) and was regulated by HAB1—a downstream component of RCAR5 in ABA signaling (Fig 4A and B). In addition, dry and imbibed *Pro35S:RCAR5* seeds showed high expression levels of ABA-responsive genes, suggesting that germination was delayed by enhanced ABA signaling (Fig 2D). Consistently, *RCAR5* expression in dry seeds was ABA dependent and decreased significantly during seed germination (Fig 1A and C). Consistent with *RCAR5*, transcripts of *RCAR1, RCAR2*, and *RCAR6* accumulated strongly in dry seeds—regardless of transcriptional difference between Col-0 and Ler ecotypes—and were downregulated during seed germination. However, the effect of gene overexpression on seed germination under cold stress conditions occurred infrequently. Our data suggest that *RCAR5* plays a major role in the maintenance of ABA-induced seed dormancy under cold stress conditions.

In contrast to the expression pattern in dry seeds, *RCAR5* expression in leaf tissues was much weaker than that of other *RCAR* gene family members and was not influenced by ABA deficiency (Fig 5C and D). Nevertheless, *RCAR5* expression was significantly upregulated in response to cold stress, whereas expression of *RCAR7, RCAR8, RCAR10, RCAR11, RCAR12*, and *RCAR13* was downregulated (Fig 5A). *RCAR5* expression was also induced via an ABA-independent pathway: Under cold-stress conditions, ABA-deficient *aba1-6* and ABA-insensitive *abi5* mutants showed similar *RCAR5* expression patterns to WT plants (Fig 5B). Moreover, *RCAR5* expression was influenced by dehydration, but not by ABA treatment (Appendix Fig S5E). Some RCARs—including RCAR1, RCAR3, and RCAR10—inhibit PP2Cs in the absence of ABA (Hao *et al.*, 2011). Hence, we predicted that *RCAR5* functions in plant cold stress responses via an ABA-independent pathway. *Pro35S:RCAR5* plants showed enhanced stomatal closure in response to cold stress (Fig 6). Our results are consistent with those of Wilkinson *et al.* (2001), who reported rapidly induced stomatal closure in the leaves of *Commelina communis* in response to cold stress, possibly owing to increased uptake of apoplastic calcium, but not ABA, into guard cells. This phenomenon seems to be specific to *Pro35S:RCAR5* plants; ABA-hypersensitive *Pro35S:RCAR2* and *Pro35S:RCAR3* mutants did not show enhanced stomatal closure in response to cold stress (Appendix Fig S6A–C). We further showed that leaf temperatures of loss-of-function mutants of *HAB1* and *PP2CA—*which can interact with RCAR5*—*did not differ significantly from those of WT plants in response to cold stress (Appendix Fig S3E-H). However, we cannot rule out the involvement of a pleiotropic effect of *Pro35S:RCAR5* plants and functional redundancy of HAB1 and PP2CA. We questioned whether cold-stress induced rapid stomatal closure in *Pro35S:RCAR5* plants occurs via an ABA-independent pathway. *Pro35S:RCAR5* plants showed ABA hypersensitivity and ABA acts as the main signal for triggering stomatal closure, hence, the involvement of ABA in the process remains unclear. Previous studies have suggested that cold stress triggers ABA biosynthesis and that ABA plays an important role in the cold stress response—in particular cold acclimation, as shown in ABA-deficient and ABA-insensitive mutants (Kushiro *et al.*, 2004, Lång *et al.*, 1994, Thomashow, 1999, Xiong *et al.*, 2001). For example, ABA-deficient mutants *aba1* and *aba3* showed impaired freezing tolerance characterized by low expression levels of ABA-responsive genes that may function in cold stress (Xiong *et al.*, 2001). Additionally, application of exogenous ABA triggers cold acclimation and enhanced freezing tolerance in Arabidopsis (Thomashow, 1999). However, ABA biosynthesis seems not to be an early event in the cold stress response. In Arabidopsis, endogenous ABA accumulated weakly at 6 h, and its level increased up to 4-fold after 15 h exposure to 4°C day/2°C night temperatures (Lång *et al.*, 1994). Moreover, expression of ABA synthesis-related genes was not upregulated at 3 h, 6 h, or 24 h after exposure to 0°C (Lee *et al.*, 2005). Here, surface temperature and fresh weight of leaves differed significantly between *Pro35S:RCAR5* and WT plants 4 h after cold treatment. In comparison with the timing of ABA biosynthesis, stomatal closure of *Pro35S:RCAR5* plants seems to start earlier. In addition, *RCAR5* overexpression in *aba1-6* mutants led to enhanced stomatal closure in response to cold stress. Our data imply that *RCAR5* is induced in response to cold stress and contributes to cold stress-induced stomatal closure via an ABA-independent pathway.

Thus, our study provides evidence for two different functions of RCAR5 via ABA-dependent and ABA-independent pathways in plant tissues such as seeds and leaves. However, the precise function of RCAR5 in the cold stress response remains unclear. Further studies to elucidate the functional interactions RCAR5–HAB1 and PP2CA–OST1 in germination delay and rapid stomatal closure under cold stress conditions, and to identify transcription factors that regulate RCAR5 expression via ABA-dependent and/or ABA-independent pathways, are required.

## Materials and Methods

### Plant material and growth conditions

Here, *Arabidopsis thaliana* (ecotype Col-0 and Ler) plants were used as the wild-type (WT). The T-DNA insertion mutants *abi5* (SALK_013163) (Zheng *et al.*, 2012a), *hab1* (SALK_002104) (Saez *et al.*, 2004b), and *pp2ca-1* (SALK_124564; Lim et al., 2014), and the EMS-mutagenized mutant *aba1-3* (Ler background) (Koornneef *et al.*, 1982) (Koornneef, Jorna et al., 1982) ^62 61^ and *aba1-6* (Col-0 background), were obtained from the Arabidopsis Biological Resource Center. Plants were routinely grown in soil mixture containing a 9:1:1 ratio of peat moss, perlite and vermiculite. The plants were maintained at 24°C and 60% humidity under fluorescent light (130 μmol photons·m^-2^·s^-1^) with a 16 h light/8 h dark cycle. Prior to in vitro culture, seeds of Arabidopsis were surface sterilized with 70% ethanol for 1 min and treated with 2% sodium hydroxide for 10 min. The seeds were then washed 10 times with sterile distilled water and sown on MS agar plates (Sigma, St. Louis, MO) supplemented with 1% sucrose. Following stratification at 4°C for 2 days, the plates were sealed and incubated at 24°C in a chamber under fluorescent light (130 μmol photons·m^-2^·s^-1^) with a 16 h light/8 h dark cycle. For tobacco (*Nicotiana benthamiana*) plants, seeds were sown in a steam-sterilized compost soil mix (peat moss, perlite, and vermiculite, 5:3:2, v/v/v), sand, and loam soil (1:1:1, v/v/v). The tobacco plants were raised in a growth chamber at 25 ± 1°C under the conditions described above.

### Generation of transgenic *RCAR*-overexpressing mutants

We previously generated Arabidopsis transgenic lines overexpressing *RCAR2* (Lee *et al.*, 2013), *RCAR3* (Lim *et al.*, 2014), *RCAR4*, and *RCAR5* (Lim & Lee, 2015). Here, we generated transgenic lines overexpressing each of the remaining *RCAR* gene family members in Arabidopsis. The coding sequences of the 10 *RCARs* were cloned into the pENTR/D-TOPO vector (Invitrogen, Carlsbad, CA, USA) and integrated into pK2GW7 using the LR reaction to induce constitutive expression of each *RCAR* under the control of the cauliflower mosaic virus 35S promoter (Karimi *et al.*, 2002). The correct construct was introduced into *Agrobacterium tumefaciens* strain GV3101 via electroporation. We conducted *Agrobacterium*-mediated transformation using the floral dip method (Clough & Bent, 1998). For selection of transgenic lines, seeds harvested from the putative transformed plants were plated on MS agar plates containing 50 μg ml^−1^ kanamycin.

### Cold treatment and phenotypic analyses

To measure the germination rate of WT and transgenic plants under cold stress conditions, seeds were vernalized at 4°C for 2 days and continuously incubated in darkness until the indicated time points. To analyze whether cold sensitivity of Arabidopsis plants is associated with ABA signaling, seeds were plated on MS agar medium supplemented with ABA (0.75 μM) and/or norflurazon (NF; 25 μM or 50 μM) as the ABA biosynthesis inhibitor.

For freezing tolerance assays, 3-week-old Arabidopsis seedlings were exposed to 4 ± 1°C for 0.5 h, followed by 1°C for 1 h. To set experimental temperatures, the temperature was incrementally decreased by 2°C over a period of 30 min and maintained for 1 h. After freezing treatment, plants were incubated at 4 ± 1°C in the dark for 12 h and then transferred to normal growth conditions. The survival rate after 4 days was measured.

For thermal imaging analysis, 2-week-old Arabidopsis seedlings were subjected to cold stress by exposure to 4 ± 1°C until the indicated time points. Thermal images were obtained using an infrared camera (FLIR systems; T420), and leaf temperatures were measured with FLIR Tools+ version 5.2 software.

### Electrolyte leakage test

Electrolyte leakage of WT and *Pro35S:RCAR5* transgenic plants subjected to freezing stress was measured as described previously, but with modifications to sample type and incubation time (Ding *et al.*, 2015). For cold acclimation, 2-week-old plants grown in soil were treated at 4 ± 1°C for 2 days and then exposed to −4°C for 1 h prior to an electrolyte leakage test. Leaf samples were detached from each plant line and placed into tubes containing 20 mL of deionized water (S0). Following shaking for 12 h at room temperature, the conductivity of the samples was measured (S1). After autoclaving for 15 min, the tubes were shaken for 1 h and the conductivity was measured (S2). Electrolyte leakage of each sample was calculated using the formula S1 – S0/S2 – S0.

### RNA isolation and quantitative reverse transcription-polymerase chain reaction

Total RNA isolation and reverse transcription-polymerase chain reaction (RT-PCR) analyses were performed as described previously (Lim & Lee, 2016). Two-week-old WT and transgenic plants were subjected to cold stress (4°C) and leaf samples were harvested at the indicated time points. cDNA was synthesized using a Transcript First Strand cDNA Synthesis kit (Roche, Indianapolis, IN) with 1 µg of total RNA according to the manufacturer’s instructions. For quantitative RT-PCR (qRT-PCR) analysis, the synthesized cDNA was amplified in a CFX96 Touch™ Real-Time PCR detection system (Bio-Rad, Hercules, CA) with iQTMSYBR Green Supermix and specific primers (Appendix Table 1). All reactions were performed in triplicate. The relative expression level of each gene was calculated using the ΔΔCt method, as described previously (Livak & Schmittgen, 2001). The Arabidopsis actin8 gene (*AtACT8*) was used for normalization.

### Stomatal aperture bioassay

The stomatal aperture bioassay was conducted as described previously, but with some modifications (Lim *et al.*, 2014). Briefly, leaf peels were collected from the rosette leaves of 3-week-old plants and floated on stomatal opening solution (SOS; 50 mM KCl, 10 mM MES-KOH, 10 μM CaCl_2_, pH 6.15). The peels were incubated at 24°C under fluorescent light for 3 h to obtain >80% stomatal opening in *Arabidopsis thaliana* Col-0 plants. The buffer was replaced with fresh SOS and leaf peels were further incubated at 4°C under fluorescent light for 3 h. In each individual sample, 100 stomata were randomly observed under a Nikon Eclipse 80i microscope. The widths and lengths of individual stomatal pores were recorded using Image J 1.46r software (http://imagej.nih.gov/ij). Each experiment was performed in triplicate.

### Yeast two-hybrid assay

Yeast two-hybrid analysis was performed as described previously (Lim & Lee, 2016). The coding sequences of RCAR5 as bait or ABI1, HAB1, AHG1, and PP2CA as prey were subcloned into pGBKT7 or pGADT7, respectively. Using the lithium acetate-mediated transformation method, these bait and prey pairs were co-transformed into yeast strain AH109. To evaluate the interaction between them, transformant candidates were screened on SC-Leu-Trp and SC-Ade-His-Leu-Trp media.

### Bimolecular fluorescence complementation assay

The bimolecular fluorescence complementation (BiFC) assay was performed as described previously (Lim & Lee, 2015). To generate the BiFC constructs, the coding sequences of *RCAR5, HAB1*, and *PP2CA* were subcloned into the *Pro35S:VYNE* and *Pro35S:CYCE* vectors using the LR reaction (Waadt *et al.*, 2008). The resulting plasmids were transformed into *Agrobacterium tumefaciens* strain GV3101. For agroinfiltration-based transient gene expression, *Agrobacterium tumefaciens* harboring *RCAR5:VYNE* combined with either *HAB1:CYCE* or *PP2CA:CYCE* were syringe-infiltrated into the leaves of 5-week-old *Nicotiana benthamiana* seedlings together with the silencing suppressor p19 in a 1:1:1 volume ratio at a final OD_600_ = 0.5 for each. At 3 days after infiltration, leaf discs were cut and the lower epidermal cells were examined under a confocal microscope (510 UV/Vis Meta; Zeiss, Oberkochen, Germany) equipped with LSM Image Browser software.

## Acknowledgments

This work was supported by a grant from the “Next-Generation BioGreen 21 Program for Agriculture & Technology Development (Project No. PJ01316801),” Rural Development Administration, and by the National Research Foundation of Korea (NRF) grant funded by the Korea Government (MSIT) (No. 2018R1A5A1023599, SRC), Republic of Korea.

## Author Contributions

C.W.L and S.C.L conceptualized and designed the study. C.W.L performed all the experiments. S.C.L supervised the project. C.W.L and S.C.L wrote the manuscript.

## Appendix Figures

**Appendix Figure 1 Expression patterns of *RCAR* genes during seed germination and enhanced ABA sensitivity of *Pro35S:RCAR* transgenic plants during seed germination and seedling growth.** (A) Relative expression levels of *RCAR* genes in dry seeds of Arabidopsis ecotype Col-0 (upper) and Ler (bottom). *Actin8* was used as an internal control for normalization and the expression level of *RCAR1* from each ecotype was set to 1.0. (B and C) Relative expression levels of *RCAR* genes during germination. Data were obtained from the Arabidopsis eFP Browser with ‘Germination’ (B) (Winter *et al.*, 2007) and ‘Klepikova Atlas’ (C) (Klepikova *et al.*, 2016) as data sources in the Bio-Analytic Resource for Plant Biology (http://bar.utoronto.ca/efp/cgi-bin/efpWeb.cgi). H, seeds from the silque; S, stratification; L, light. (D) Expression levels of *RCAR* genes in the leaves of *Pro35S:RCAR* transgenic plants. *Actin8* was used as an internal control for normalization and the expression level of each *RCAR* gene in WT plants was set to 1.0. (E–G) Seedling development of *Pro35S:RCAR* transgenic lines and WT plants in the presence of ABA. Seeds of *Pro35S:RCAR* transgenic lines were germinated on 0.5× MS medium supplemented with 0 μM or 0.5 μM ABA and grown at 24°C in the light. At 5 days after incubation (DAI), cotyledon greening (F) and root length (G) were measured and representative images were taken (E). Data represent mean ± SD of three independent experiments.

**Appendix Figure 2 ABA sensitivity of *aba1-6, abi5, Pro35S:RCAR5/aba1-6*, and *Pro35S:RCAR5/abi5* transgenic plants during seed germination and seedling growth.** (A) Expression levels of *RCAR* genes in the leaves of *Pro35S:RCAR/aba1-6* transgenic plants. *Actin8* was used as an internal control for normalization and the expression level of *RCAR5* in WT plants was set to 1.0. (B) Germination rates of *Pro35S:RCAR, Pro35S:RCAR/aba1-6, aba1-6*, and WT plants on 0.5× MS medium supplemented with 0 μM or 0.5 μM ABA. The numbers of seeds with emerged radicles were counted 3 days after plating. (C–E) Seedling development of *Pro35S:RCAR, Pro35S:RCAR/aba1-6, aba1-6*, and WT plants in the presence of ABA. Seeds of each plant line were germinated on 0.5× MS medium supplemented with 0 μM, 0.5 μM, or 0.75 μM ABA and vertically grown at 24°C in the light. At 7 days after incubation (DAI), root length (D) and cotyledon greening (E) were measured and representative images were taken (C). (F) Expression levels of *RCAR* genes in the leaves of *Pro35S:RCAR/abi5* transgenic plants. *Actin8* was used as an internal control for normalization and expression level of *RCAR5* in WT was set to 1.0. (G) Germination rates of *Pro35S:RCAR, Pro35S:RCAR/abi5, abi5*, and WT plants on 0.5× MS medium supplemented with 0 μM or 0.5 μM ABA. The numbers of seeds with emerged radicles were counted 3 days after plating. (H–J) Seedling development of *Pro35S:RCAR, Pro35S:RCAR/abi5, abi5*, and WT plants in the presence of ABA. Seeds of each plant line were germinated on 0.5× MS medium supplemented with 0 μM or 0.75 μM ABA and vertically grown at 24°C in the light. At 7 DAI, root length (I) and cotyledon greening (J) were measured and representative images were taken (H). Data represent mean ± SD of three independent experiments. Different letters indicate significant differences between three independent experiments (ANOVA; *P* < 0.05). Scale bar = 1 cm.

**Appendix Figure 3 Expression patterns of PP2C genes during seed germination and after cold stress treatment, and reduced ABA sensitivity of *Pro35S:HAB1* and *Pro35S:RCAR5/HAB1* transgenic plants during seed germination and seedling growth.** (A and B) Relative expression levels of *PP2C* genes during germination. Data were obtained from the Arabidopsis eFP Browser with ‘Germination’ (A) (Winter *et al.*, 2007) and ‘Klepikova Atlas’ (B) (Klepikova *et al.*, 2016) as data sources in the Bio-Analytic Resource for Plant Biology (http://bar.utoronto.ca/efp/cgi-bin/efpWeb.cgi). H, seeds from the silque; S, stratification; L, light. (C and D) Relative expression levels of *PP2C* genes in Arabidopsis shoots and roots in response to cold stress. Data were obtained from the Arabidopsis eFP Browser with ‘Abiotic stress’ (Winter *et al.*, 2007) as the data source. (E) Expression levels of *RCAR* genes in the leaves of *Pro35S:HAB1* and *Pro35S:RCAR/Pro35S:HAB1* transgenic plants. *Actin8* was used as an internal control for normalization. The expression levels of *RCAR5* or *HAB1* genes in WT plants were set to 1.0. (F) Germination rates of *Pro35S:RCAR, Pro35S:HAB1, Pro35S:RCAR/Pro35S:HAB1*, and WT plants on 0.5× MS medium supplemented with 0 μM, 0.5 μM, or 1 μM ABA. The numbers of seeds with emerged radicles were counted 3 days after plating. (G–I) Seedling development of *Pro35S:RCAR, Pro35S:HAB1, Pro35S:RCAR/Pro35S:HAB1*, and WT plants in the presence of ABA. Seeds of each plant line were germinated on 0.5× MS medium supplemented with 0 μM, 0.5 μM, or 0.75 μM ABA and vertically grown at 24°C in the light. At 7 DAI, root length (H) and cotyledon greening (I) were measured and representative images were taken (G). Data represent mean ± SD of three independent experiments. Different letters indicate significant differences between three independent experiments (ANOVA; *P* < 0.05). Scale bar = 1 cm.

**Appendix Figure 4 Sensitivity of *hab1* and *pp2ca* mutant plants to cold stress.** (A) Physical interaction of RCAR5 with HAB1 and PP2CA in yeast. Interaction was indicated by growth on selection medium (SC-Ade-His-Leu-Trp); growth on SC-Leu-Trp was used as a control (right). (B) BiFC assay of interactions of RCAR5 with HAB1 and PP2CA. RCAR5:VYNE was co-expressed with HAB1:CYCE or PP2CA:CYCE in the leaves of tobacco. Scale bar = 20 mm. (C) Germination rates of *Pro35S:RCAR5, hab1, pp2ca-1*, and WT plants. Seeds of these plants were plated on 0.5× MS medium and germinated in the dark at 4°C. (D) Vertical growth of *Pro35S:RCAR5, hab1, pp2ca-1*, and WT plants exposed to ABA or LT. Seeds of each plant were plated on 0.5× MS medium supplemented with 0 μM or 0.75 μM ABA and germinated under normal or cold stress conditions. Scale bar = 1 cm. Representative images were taken at 8 DAI (for ABA) and 18 DAI (for LT). (E) qRT-PCR analysis of *HAB1* and *PP2CA* gene expression in Arabidopsis leaves in response to cold stress (4°C). *Actin8* was used as an internal control for normalization. The expression level of each gene at 0 h was set to 1.0. (F–H) Leaf dehydration and leaf temperatures of *Pro35S:RCAR5, hab1, pp2ca-1*, and WT plants in response to cold stress. Representative thermographic images of *Pro35S:RCAR5, hab1, pp2ca-1*, and WT plants 24 h after exposure to 4°C (F); the mean leaf temperatures of the two largest leaves were measured using 20 plants of each line (G). Concomitantly, the fresh weights of each plant line were measured (H). All data represent mean ± SD of three independent experiments. Different letters indicate significant difference between WT and transgenic plants (ANOVA; *P* < 0.05).

**Appendix Figure 5 Expression patterns of *RCAR* genes in Arabidopsis leaves after cold stress treatment.** (A) Relative expression levels of *RCAR* genes in the leaves of Arabidopsis ecotype Col-0. *Actin8* was used as an internal control for normalization, and the expression level of *RCAR1* was set to 1.0. (B) Relative expression levels of *RCAR* genes in the leaves of *aba1-6* and WT plants. (C–D) Relative expression levels of *PP2C* genes in Arabidopsis shoots and roots in response to cold stress. Data were obtained from the Arabidopsis eFP Browser with ‘Abiotic stress’ (Winter *et al.*, 2007) as the data source in the Bio-Analytic Resource for Plant Biology (http://bar.utoronto.ca/efp/cgi-bin/efpWeb.cgi). (E) Expression patterns of *RCAR2, RCAR3*, and *RCAR5* in Arabidopsis leaves after dehydration and ABA treatment. Relative expression levels of *RCAR* genes in Arabidopsis leaves after ABA treatment and dehydration stress. *Actin8* was used as an internal control for normalization. The expression level of each gene at 0 h was set to 1.0. (F and G) Enhanced dehydration tolerance of *Pro35S:RCAR5* transgenic plants. (F) Dehydration sensitivity of *Pro35S:RCAR5* and WT plants. Plants were subjected to dehydration stress by withholding water for 12 days (n = 16); representative images were taken 3 days after rewatering. (G) Water loss from leaves of WT and transgenic plants at various times after detachment of leaves. Data represent mean ± standard error of three independent experiments (n = 20). Different letters indicate significant difference between WT and transgenic plants (ANOVA; *P* < 0.05).

**Appendix Figure 6 Cold stress-induced dehydration of WT and *Pro35S:RCAR2, Pro35S:RCAR3*, and *Pro35S:RCAR5* transgenic plants.** (A and B) Phenotypic response of WT and transgenic lines to LT. Three-week-old Arabidopsis plants were exposed to 4°C for 24 h and images were taken (A). Representative parts were magnified (B). Arrow heads indicated wilted leaves. (C) Water loss from WT and *Pro35S:RCAR2, Pro35S:RCAR3*, and *Pro35S:RCAR5* plants after cold stress treatment. The fresh weights of each plant line were measured 24 h after treatment. Data represent the mean ± SE of three independent experiments, each evaluating 30 plants. Different letters indicate significant difference between WT and transgenic plants (ANOVA; *P* < 0.05). (D and E) Thermal imaging analysis of *Pro35S:RCAR5* transgenic plants. Representative thermographic images of WT and *Pro35S:RCAR5* plants after LT treatment. Thermographic images were taken at different time points after LT treatment (D), and the mean leaf temperatures of the two largest leaves were measured using 20 plants of each line (E). Different letters indicate significant differences (ANOVA; *P* < 0.05).

**Appendix Figure 7 Strong upregulation of cold- and ABA-responsive genes in *Pro35S:RCAR5 Pro35S:RCAR/Pro35S:HAB1*, and *Pro35S:RCAR/abi5* plants under cold stress conditions.** Two-week-old *Pro35S:RCAR5* (A), *Pro35S:RCAR/Pro35S:HAB1* (B), *Pro35S:RCAR/abi5* (C) and WT plants were exposed to 4°C and shoots were harvested at the indicated time points. *Actin8* was used as an internal control for normalization. The expression level of each gene at 0 h was set to 1.0. Data represent mean ± SD of three independent experiments. Different letters indicate significant difference between WT and transgenic plants (ANOVA; *P* < 0.05).

**Appendix Table 1** Sequences of primers used in this study.

